# Contaminant Spot Check and Removal Assay (ContamSPOT) for Mass Spectrometry Analysis

**DOI:** 10.1101/2024.01.05.574392

**Authors:** Noah Smeriglio, Haorong Li, Wan Nur Atiqah binti Mazli, Katharine Bendel, Ling Hao

**Author notes:** Corresponding author Ling Hao, Assistant Professor Department of Chemistry, Department of Biochemistry and Molecular Medicine George Washington University.

## Abstract

Mass spectrometry (MS) analysis is often challenged by contaminations from detergents, salts, and polymers that compromise the data quality and can damage the chromatography and mass spectrometry instruments. However, researchers may not discover contamination issues until after data acquisition. There is no existing contaminant assay that is sensitive enough to detect trace amounts of contaminants from a few microliters of samples prior to MS analysis. To address this crucial need in the field, we developed a sensitive, rapid, and low-cost contaminant spot check and removal assay (ContamSPOT) to detect and quantify trace amounts of contaminants, such as detergents, salts, and other chemicals commonly used in MS sample preparation workflow. Only one microliter of sample was used prior to MS injection to quantify contaminants by ContamSPOT colorimetric or fluorometric assay on a thin layer chromatography (TLC) plate. We also optimized contaminant removal methods to salvage samples with minimal loss when ContamSPOT showed a positive result. ContamSPOT was then successfully applied to evaluate commonly used bottom- up proteomic methods, regarding the effectiveness of removing detergents, peptide recovery, and reproducibility. We expect ContamSPOT to be widely adopted by MS laboratories as a last-step quality checkpoint prior to MS injection. ContamSPOT can also provide a unique readout of sample cleanness for developing and evaluating MS-based sample preparation methods. We provided a practical decision tree and a step-by-step protocol with a troubleshooting guide to facilitate the use of ContamSPOT by other researchers.

**Table of Contents (TOC) graphic:** (All images used in the TOC graphic are original, in compliance with copyright guidelines.)

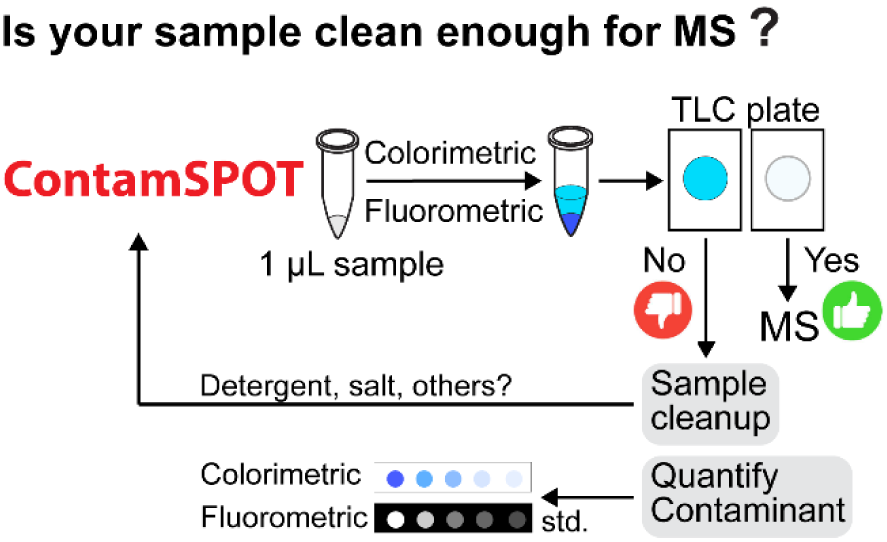

## INTRODUCTION

Minimizing contaminations prior to mass spectrometric (MS) analysis has been a long-standing challenge for analytical chemists.^1–4^ Contaminants such as detergents, polymers, solvents, and salts can significantly influence the separation and ionization efficiency of target molecules in chromatography and MS.^5^ Trace amount of detergent contaminations (SDS, Triton, Tween, *etc.*) were notoriously known to strip packing materials from chromatography columns and damage both the chromatography and MS instruments.^6,7^ Extensive contaminations can harm the ion funnel and mass analyzer beyond the ion source, leading to costly downtime for deep cleaning and repair services. Contaminants are introduced throughout the experimental workflow and may not be discovered until after data acquisition, already damaging the instrument and losing precious samples. Contaminations can be challenging to identify when samples were collected from another lab for a collaborative project or received by an MS facility. Therefore, developing a fast quality checkpoint for contaminants prior to MS analysis can greatly benefit the researchers and protect valuable samples and expensive instruments.

Detergents and salts are often added to buffers to facilitate the extraction of membrane proteins or lipids, which need to be removed in downstream sample preparation steps. Various sample preparation workflows have been developed for detergent-containing samples or removing contaminants prior to MS analysis for different types of molecules.^7–13^ For bottom-up proteomics, filter-based FASP^14^ and S-Trap^15^ methods can remove contaminants prior to protein digestion. Recently developed beads-based SP3^16^, SP2^17^ and SP4^18^ methods can remove contaminants at the protein level or peptide level. However, the efficiency of removing contaminants with these methods can only be assessed by examining the MS data due to the lack of a sensitive and quantitative contaminant assay. Inadequate sample cleanup compromises data quality and harms the instrument, while excessive cleanup causes sample loss and increases both the time and cost of sample preparation. Therefore, a sensitive and quantitative contaminant assay can provide a unique readout of sample cleanness for method development and decision-making regarding sample cleanup.

A practical contaminant assay prior to MS analysis should be rapid, highly sensitive, and require minimal sample volume. Detergents such as SDS have been traditionally quantified by spectroscopy methods in environmental analysis.^19,20^ However, these methods typically require a minimal volume of 20-100 μL, and do not provide sufficient sensitivity to quantify the trace amount of contaminants in the sample after routine cleanup steps. Ideally, the method should also be straightforward and low-cost, so that researchers are willing to incorporate this assay as a last-step quality checkpoint prior to MS analysis.

To address this crucial need in the field, we developed a Contaminant Spot Check and Removal Assay (ContamSPOT) that only consumes 1 μL of sample within a few minutes prior to MS analysis. We screened various solvents and reagents to extract common MS contaminants and amplify their signals, so that they could be spotted and quantified on a thin layer chromatography (TLC) plate. We developed both colorimetric and fluorometric assays to expand the applicability of ContamSPOT to different types of contaminants, including ionic and non-ionic detergents, salts, acids, and other chemicals. We then optimized contaminant removal methods so that contaminants can be effectively removed with minimal sample loss when ContamSPOT showed positive results. We then applied ContamSPOT to compare several popular proteomics sample preparation methods that should remove contaminants at the protein or peptide level. Lastly, we provided a practical decision tree and a step-by-step protocol to assist users in conducting ContamSPOT assay and making experimental decisions based on ContamSPOT results.

## EXPERIMENTAL METHODS

### ContamSPOT Colorimetric and Fluorometric Assays

For the colorimetric assay, a mixture was created in a 0.2 mL tube containing 1 µL of sample, 1 µL of ion-pairing agent (0.1% o-toluidine blue, Carolina Biological) and 3 µL of ethyl acetate (ACS grade, Fisher Scientific). The mixture was vortexed and centrifuged briefly. One to two microliters of the top layer were spotted onto a TLC plate (Sorbtech). The spot dried instantly and the blue color (indicating presence of contaminants) can be directly visualized by eye. An image can be taken and quantified by ImageJ software.^21^ To accurately quantify the amount of contaminant from an unknown sample, calibration curve samples using series concentrations of contaminant standards should be spotted on the same TLC plate with the unknown sample.

For the fluorometric assay, reagents were obtained from the ProFoldin Detergent Assay Kit (DAK 1000) with a modified protocol: A mixture was created in a 0.2 mL tube containing 1 µL of sample, 2 µL of 1X ProFoldin dye A43, and 1 µL of 1X Reagent 2. The mixture was vortexed briefly then incubated at room temperature for 5 min. Two microliters of the mixture were spotted onto a TLC plate (no visible color). A Fluorescence Imager (Bio-Rad ChemiDoc^TM^ MP) was used to image the plate at 535 nm, followed by quantification in ImageJ. Detailed step-by-step protocol is provided in **Supporting Information**.

### Ethyl Acetate Liquid-Liquid Extraction to Remove Detergents

Water-saturated ethyl acetate was created by mixing pure ethyl acetate (HPLC grade, Sigma) with a small amount of HPLC grade water in a glass bottle until forming water layer at the bottom. Water-saturated ethyl acetate was added at 10-fold volume to the samples, followed by 30 s of vortex and 30 s of centrifugation. Then the top ethyl acetate layer was carefully removed without touching the bottom aqueous layer. This process was repeated for a total of 3-5 extraction cycles. Cleaned samples were then dried and ready for LC-MS analysis. See **Supporting Information** for detailed steps.

### Cell Culture and Protein Extraction

HEK293 cells were routinely cultured and harvested in the lab as described previously.^4,22^ Cell pellets were lysed in either urea buffer (8M urea, 150 mM NaCl, 50 mM ammonium bicarbonate (AmBC)) or detergent buffer (2% SDS, 1% Triton X-100, 150 mM NaCl, 50 mM AmBC), followed by sonication, clarification by centrifugation, and protein concentration measurement. Protein lysates were routinely reduced and alkylated by TCEP, IAA, and DTT treatments as described previously. ^4^

### Protein Sample Preparation using S-Trap, SP3, SP4, and Acetone Precipitation Methods

Reduced and alkylated protein lysates in detergent buffer were aliquoted and prepared using several commonly used methods that can remove contaminations at the protein level, followed by trypsin digestion. Tryptic peptides from various methods were acidified by 1% formic acid (FA), dried and stored in -30 °C. S-Trap sample preparation followed Protifi manufacturer protocol with minor adjustments.^15^ Briefly, protein lysate was diluted with 100 mM triethylammonium bicarbonate (TEAB) in 90% methanol to 200 μL and applied to Micro S-trap columns. After three TEAB washes, trypsin (Promega, sequencing grade) was added at a 1:10 (trypsin: protein, w:w) ratio for digestion in 50 mM AmBC buffer at 47 °C for 2 h. Peptides were eluted using three washes: 50 mM AmBC, 0.1% FA in 2% acetonitrile (ACN) and finally 50% ACN, each centrifuged at 1,000g. SP3 sample preparation followed published protocol with minor adjustments.^16^ Briefly, protein aggregation was induced onto the Cytiva paramagnetic beads by first adding ACN to 80% to the protein lysate, followed by beads addition with a bead-to-protein ratio of 1:25 (w/w). After 15 minutes of incubation, the beads were washed twice with 95% ACN and then twice with 70% ethanol. On-bead digestion was conducted in 50 mM AmBC using a 1:30 trypsin to protein ratio for 16 h at 37 °C, followed by peptide elution on a magnetic rack. A recently developed SP4 method was conducted using glass beads enrichment (10:1, beads: protein, w/w) at 80% ACN.^18^ Sample-bead mixtures were centrifuged for 5 min at 16,000 g. The protein-bead pellet was then washed three times with 80% ethanol and resuspended in 50 mM AmBC. Samples were sonicated for 5 min, followed by trypsin digestion (1:30 ratio, 16 h), and peptide elution by centrifugation. For acetone precipitation, four-fold volume of cold acetone (-20 °C) was added to the protein lysate and placed into a -80 °C freezer overnight. Samples were then centrifuged at 15,000g to precipitate proteins at 4 °C and washed with 1 mL of cold acetone for three times. Samples were then air dried and reconstituted in 50 mM AmBC and 15 mM NaCl for trypsin digestion and desalting.

### Peptide Sample Preparation Using SP2, Detergent removal resin, and Desalting Methods

Reduced and alkylated protein lysates in urea buffer were digested overnight using 1:30 (w/w) trypsin and quenched by FA until pH <3. Known amounts of detergents were added to the peptide samples followed by several peptide cleanup methods. For SP2 cleanup, peptide samples were dried down and reconstituted in 95% ACN with a 1:25 (w/w) beads-protein ratio (Cytiva).^17^ Beads were bound to the peptides for 15 minutes followed by four 95% ACN washes. Peptides were then eluted twice into HPLC grade water. For detergent removal resin, Thermo Scientific HiPPR^TM^ detergent removal spin column kit was used here. The resin was added to the spin column and equilibrated by 50 mM AmBC three times via centrifuge at 1,500g. Samples were then added to the spin column, incubated for 2 min, and centrifuged for 1 min to elute peptides prior to acidification and desalting. For reverse-phase desalting, Oasis HLB desalting plate (Waters) was used following the manufacturer’s protocol. All samples from different protein and peptide cleanup methods were dried and reconstituted in 2% ACN and 0.1% FA for ContamSPOT, colorimetric peptide assay (Pierce), and LC-MS analysis.

### LC-MS Analysis

LC-MS/MS analysis was conducted on a Dionex Ultimate 3000 nanoLC system coupled with a Thermo Scientific Q-Exactive HFX MS. A two-hour LC gradient was used with 0.1% FA in water as buffer A and 0.1% FA in ACN as buffer B. Samples were first loaded onto a PepMap 100 C18 trap column (3 μm, 100 Å, 75 μm x 2 cm) at 5 μL/min and then separated on an Easy-spray PepMap C18 column (3 μm, 100 Å, 75 μm x 15 cm) at 0.3 μL/min and 55 °C. MS1 scanned from *m/z* 400 to 1500 with a resolving power of 60,000. A top 25 data-dependent acquisition was conducted with an MS/MS resolving power of 15,000, an isolation window of 1.4 Da, a collision energy of 30%, and a dynamic exclusion time of 22.5 s. The maximum injection times were 30 ms and 35 ms for MS and MS/MS, respectively. Automatic gain controls were set to 10^6^ and 10^4^ for MS and MS/MS, respectively.

### Proteomics Data Analysis

Proteomics data were analyzed with the Thermo Fisher Proteome Discoverer (PD, 2.4) software. Swiss-Prot *Homo sapiens* database (reviewed, 20,428 entries) and our custom-made cell culture-specific protein contaminant FASTA library (https://github.com/HaoGroup-ProtContLib)^4^ were included for protein identification. Contaminant proteins (marked with Cont_ prefix in our custom contaminant FASTA) were selected and removed from the dataset as described in our previous publication.^4^ False discovery rate cutoff was set as 1% for protein and peptide spectral matches. Maximum missed cleavages were set to 2 with trypsin as the enzyme. Precursor tolerance was set to 20 ppm. Modifications include cysteine carbamidomethylation (fixed), methionine oxidation (variable) and protein level N-terminus acetylation (variable). Further data analysis was conducted in R-studio.

## RESULTS AND DISCUSSION

### Developing a ContamSPOT Colorimetric Assay to Detect and Quantify Common MS Contaminants

We aimed to develop a fast and sensitive contaminant detection and quantification method with minimum sample consumption for MS samples. We started with the notorious SDS detergent that was often added in protein lysis buffer but not sufficiently removed before MS. A cationic dye O-toluidine blue (OTB) has been reported to form an ion-pair with SDS (**Figure 1A**), which can be used to quantify SDS from environmental samples (*e.g.* river water) using a spectrophotometer.^20^ We first screened immiscible organic solvents that can extract SDS-OTB ion-pairs to a separate layer, allowing the extraction of trace level of SDS from minimal sample volume. Six organic solvents were tested as shown in **Figure 1B**. Ethyl acetate successfully extracted SDS-OTB ion-pair into the upper organic layer, and the control sample without SDS had no color in the upper layer. At such a low volume, drying down the organic layer and redissolving into water for spectroscopy analysis did not provide sufficient sensitivity and reproducibility. Therefore, we directly spotted 1-2 μL of the top ethyl acetate layer onto a TLC plate. The spot dried instantly, and the blue color can be directly visualized by eye, indicating the presence of SDS in the sample.

**Figure 1:**
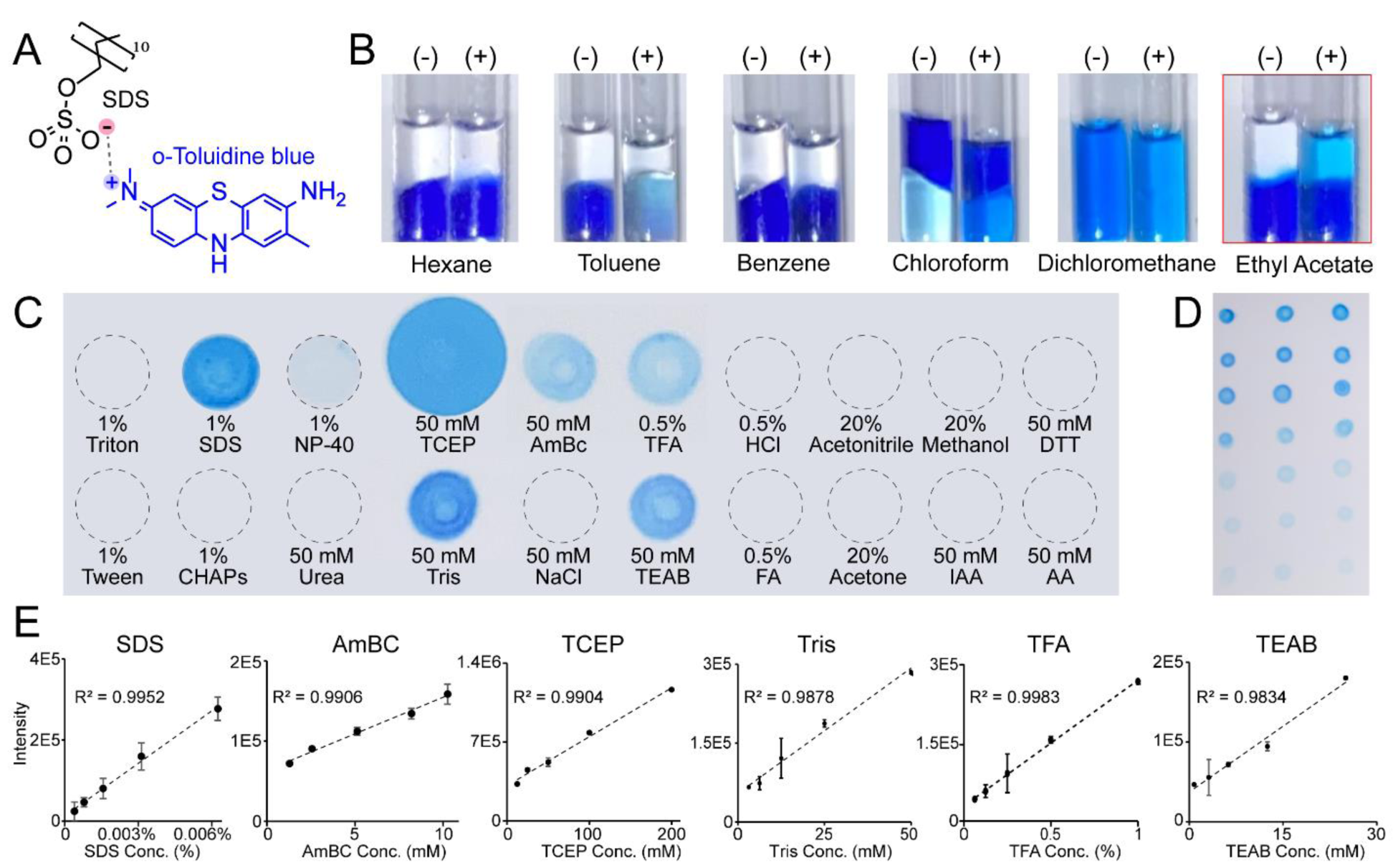
Development of the ContamSPOT colorimetric assay. (A) Schematic of the o-toluidine blue (OTB)-SDS ion-pair. (B) Solvent screening to extract the OTB-SDS ion-pair by mixing 2 μL of solvent with 1 μL of OTB dye and 1 μL of 0.1% SDS (+), compared to water control (-) in thin glass capillary tubes. (C) Screening of commonly used sample preparation chemicals. Ethyl acetate and OTB was added to each sample followed by brief vortex and centrifugation. Then 2 μL of the ethyl acetate top layer was spotted on a TLC plate. (D) Example image of a series concentration of SDS solutions with 3 technical replicates. (E) Calibration curves of positively tested chemicals.

We then tested if this method could be used to detect other contaminants besides SDS. **Figure 1C** showed the screening results of common chemicals used in MS-based sample preparation workflows. SDS, TCEP, Tris, AmBC, TFA, and TEAB showed positive blue color. To test the quantification performance of our ContamSPOT assay, we used standard solutions with series concentrations, and quantified the blue color on the imaged TLC plate by Image J software (**Figure 1D**). We then established calibration curves for all positive chemicals (**Figure 1E**). Excellent linearity and reproducibility can be obtained for all positive chemicals using ContamSPOT colorimetric assay. We also tested mixtures of several contaminants and obtained good linearity with a serial dilution. Although ContamSPOT is not selective to each tested chemical, their different performances can provide clues for identification. For example, TCEP spot was significantly wider than other chemicals. Salts like AmBC, TEAB and Tris showed pink color in the ethyl acetate layer during extraction, but the spot returned back to blue after TLC plate was air-dried. FA can be used to quench protein digestion instead of TFA in proteomics steps to avoid the interference of TFA in ContamSPOT assay. In a real experiment, researchers have prior knowledge of what chemicals were used in sample preparation steps. The sample should have been cleaned up to remove the majority of contaminants before the ContamSPOT assay. With a successful desalting, most salts should be undetectable (mM range of LODs). But trace amount of SDS can be detected and quantified from the sample with an extremely high sensitivity (LOD of 0.0004%).

### Expanding ContamSPOT with a Fluorometric Assay for Non-Ionic Detergent Contaminants

ContamSPOT colorimetric assay is limited to ionic species that can form an ion pair with OTB dye and be extracted into ethyl acetate. To expand the applicability of this assay, particularly for non-ionic detergents, such as Triton, Tween, and NP-40, we expanded the ContamSPOT method with a fluorometric assay. We adopted the A43 dye from ProFoldin’s Kit, which was used for spectroscopy analysis of non-ionic detergents. As shown in **Figure 2A**, Triton-Dye complex exhibited a shifted fluorescence wavelength away from Triton standard and peptide signals (BSA digest) with a more than 10-fold increase of signal intensity compared to Triton alone. We tested various detergents by mixing 1 uL of detergent standard with ProFoldin A43 dye, spotting 2 µL of the mixture on a TLC plate, and imaging under a fluorescence imager. As expected, non-ionic detergents Triton, CHAPS, NP-40, and Tween showed positive signals, while ionic detergent SDS and water hadno signal (**Figure 2B**). Then we established calibration curves for all positive detergents (**Figure 2C**). Fluorescence signals saturated at high concentrations, leading to nonlinear calibration curves in various fluorescence emission times and concentration ranges that we tested (**Supplemental Figure S1**). Therefore, a logarithmic function was used for curve fitting. Excellent reproducibility and sensitivity were achieved by ContamSPOT fluorometric assay with 0.002% LOD for Triton when using a longer emission capture time. In summary, we have tested a total of 22 commonly used chemicals in MS workflow, 11 of which showed positive results in ContamSPOT colorimetric or fluorometric assay, summarized in **Table 1**.

**Figure 2:**
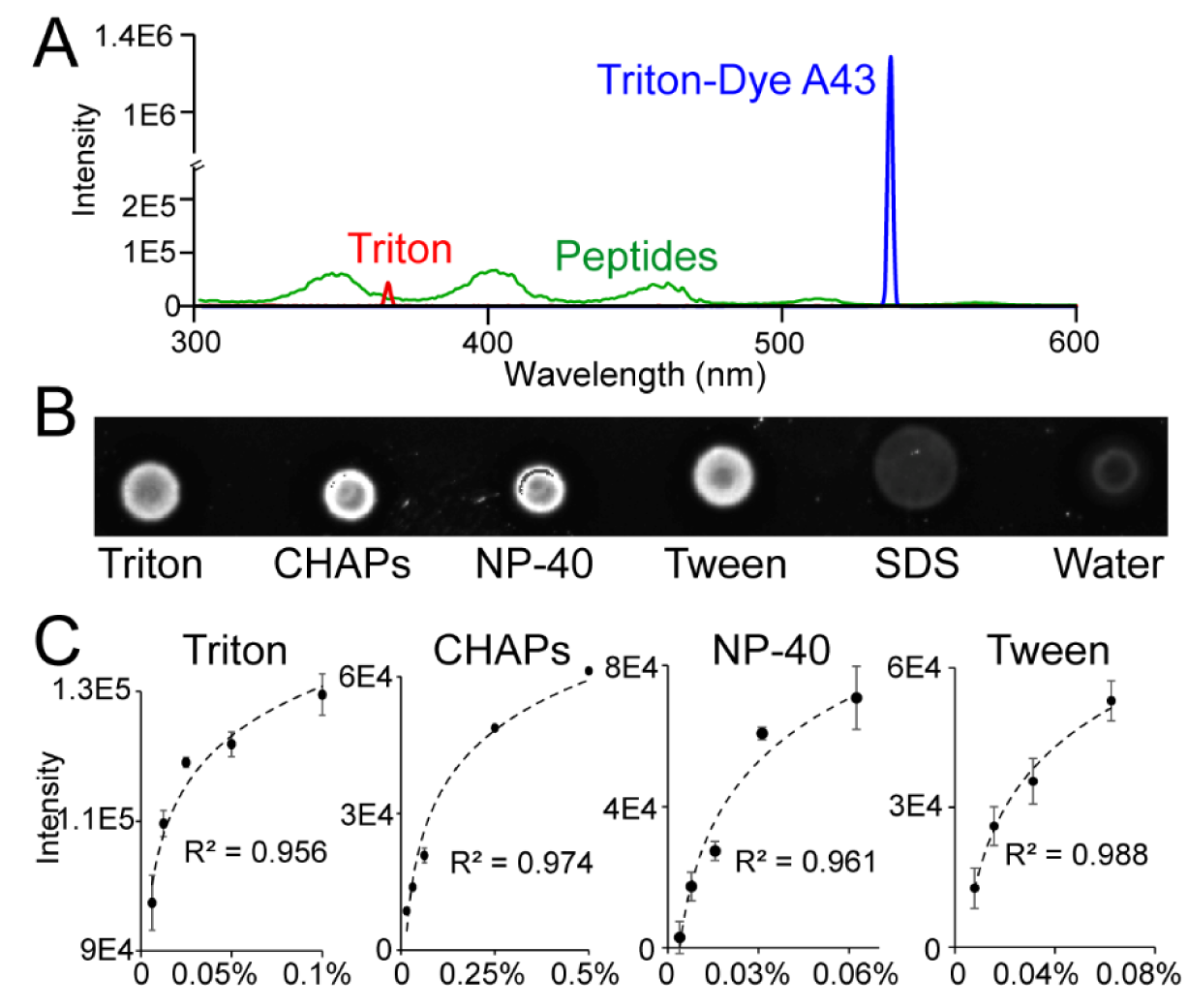
Development of the ContamSPOT fluorometric assay. (A) Fluorescence spectra of 0.1% Triton, 0.1% Triton-Dye A43 complex, and peptides from BSA digest. (B) Contaminant screening of common detergents used in lysis buffer. 1% concentration was used for each detergent. (C) Nonlinear calibration curves for positively tested detergents in ContamSPOT fluorometric assay.

**Table 1:**
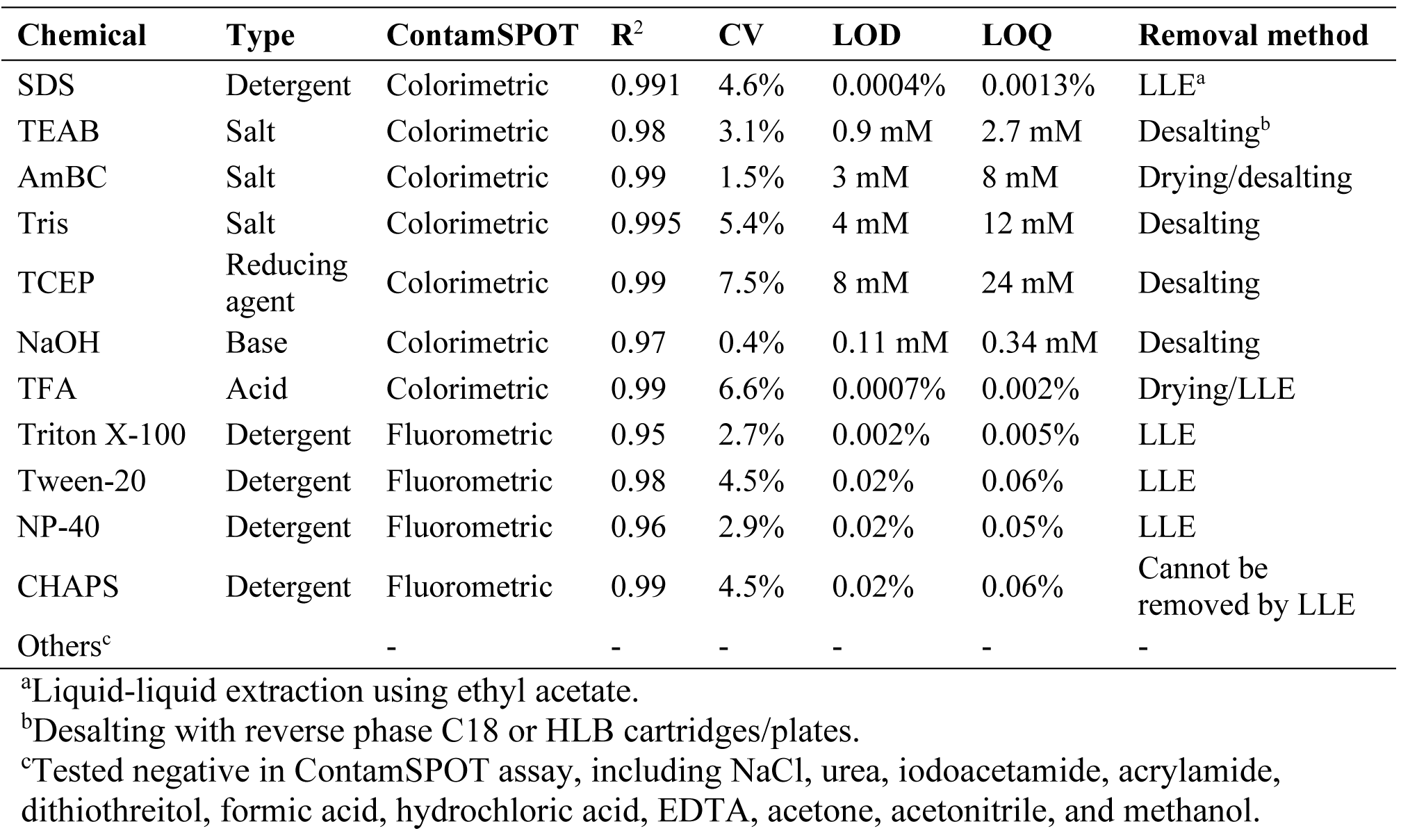
Summary of chemicals that can be detected by ContamSPOT assay.

### Optimizing Contaminant Removal Methods Prior to LC-MS Analysis

Our ContamSPOT colorimetric and fluorometric assays enabled a fast detection and quantification of various contaminants prior to MS analysis. If ContamSPOT showed positive result, we need to further clean up the sample to remove contaminants prior to MS injection. Liquid-liquid extraction (LLE) using ethyl acetate has been successfully used to remove detergents such as SDS, NP-40, Triton, OG, and DM from proteomic samples.^23,24^ Here, we further optimized the extraction experiment and screened various contaminants, enabled by our ContamSPOT colorimetric and fluorometric assays. As shown in **Figure 3A**, SDS, Triton, NP-40, and Tween can be extracted by ethyl acetate from water solution. TFA can also be extracted by ethyl acetate. However, CHAPS and other salts cannot be removed by LLE. We then optimized the LLE conditions for sufficient removal of these contaminants and minimized sample loss (**Figure 3B**). ContamSPOT assay and Pierce Peptide Colorimetric Assay were used as readouts for contaminant and peptide concentrations. Ten-fold volume of ethyl acetate can effectively remove trace amount of SDS after three ethyl acetate wash cycles with a 99% peptide recovery. If 5-fold volume is used, eight washes are needed to completely remove SDS with minimal sample loss (**Supplemental Figure S2**). Non-ionic detergents such as Triton showed high affinity to ethyl acetate and were undetectable after a single wash (**Figure 3C**). Because ethyl acetate can dissolve up to 3% water and concentrate the peptide sample during LLE, peptide recovery showed over 100% in some samples. To minimize water/peptide loss during LLE, water-saturated ethyl acetate can be used for LLE (**Figure 3E**).

**Figure 3:**
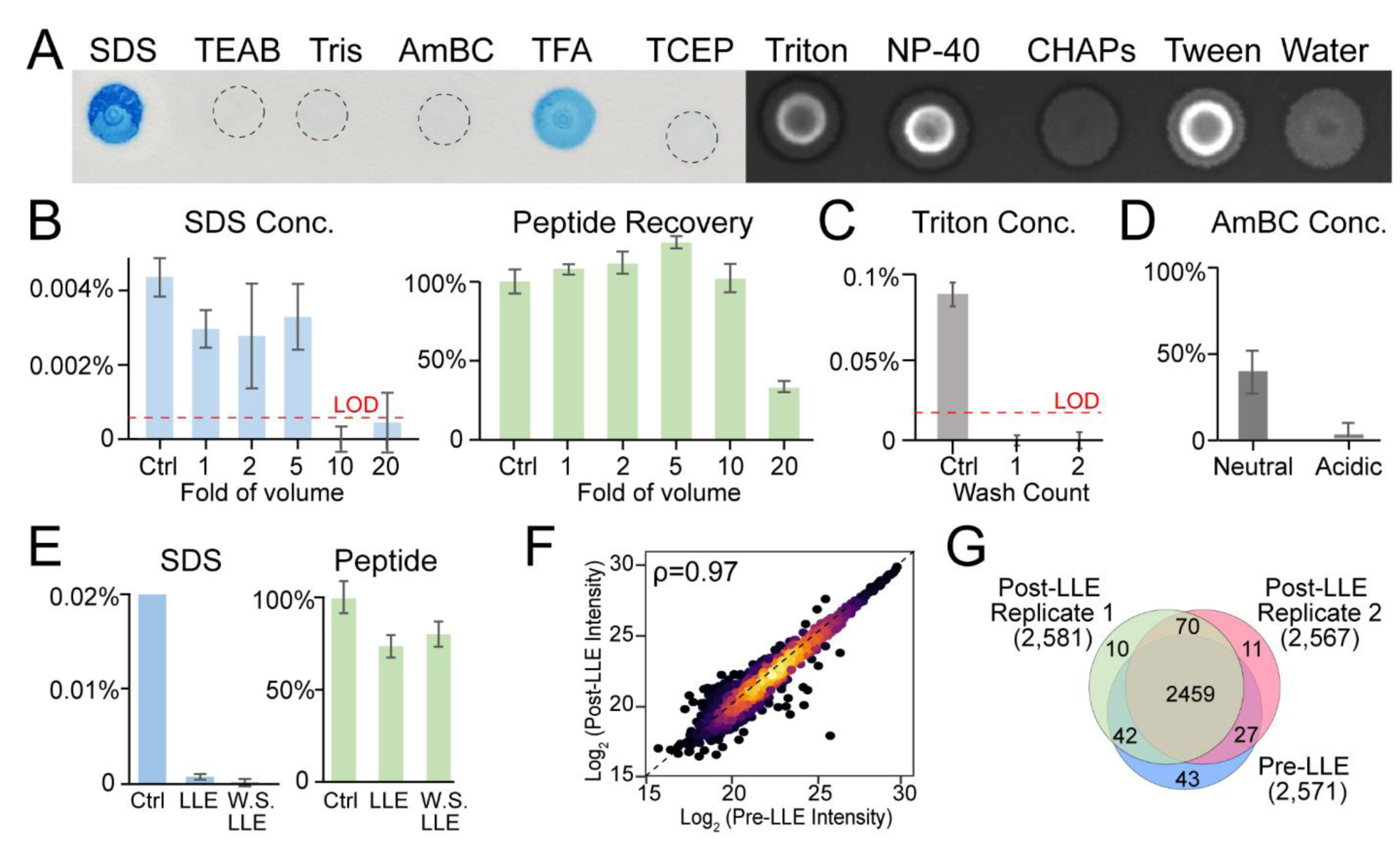
Optimization of ethyl acetate liquid-liquid extraction (LLE) method to remove contaminants. (A) ContamSPOT colorimetric (left) and fluorescence (right) assay screening of the extractants from ethyl acetate LLE. (B) SDS concentrations and peptide recoveries using different amount of ethyl acetate (fold of volume to samples) with three wash cycles. (C) Triton concentration after multiple wash cycles with 10-fold ethyl acetate volume. (D) Drying down under acidic condition improved the removal of volatile salt ammonium bicarbonate. (E) SDS concentrations and peptide recoveries after pure ethyl acetate LLE and water saturated (W.S.) ethyl acetate. (F) Scatter plot showing peptide intensity correlation before and after LLE cleanup. (G) Venn diagram of number of quantified proteins before and after LLE cleanup with two technical replicates.

To further evaluate the influence of ethyl acetate LLE on LC-MS results, we analyzed cleaned HEK peptides before and after LLE with two technical replicates. Peptide abundances showed excellent linear correlation before and after LLE with similar IDs and excellent LLE reproducibility (**Figure 3F, 3G**). The ethyl acetate extractant showed no peptide signals in LC-MS (**Supplemental Figure S3**). We did notice that ACS grade ethyl acetate extractant contains several singly charged contaminant peaks compared to HPLC grade ethyl acetate and therefore suggest using the HPLC grade ethyl acetate for LLE experiment. To summarize, ethyl acetate LLE can be used as a rescue method for detergent-contaminated samples without influencing peptide recovery and LC-MS analysis. Volatile salts can be removed by drying down, and we found that drying down under acidic condition can dramatically improve the removal of AmBC from samples (**Figure 3D**). Other nonvolatile salts have to be removed by reverse phase C18 or HLB desalting. The methods to remove different types of contaminants were summarized in **Table 1**.

### Evaluating Different Proteomic Sample Preparation Methods using ContamSPOT Assay

Various proteomic methods have been developed in recent years to prepare detergent-containing samples and remove various contaminants prior to MS analysis.^14–18^ However, the efficiency of removing detergents has not been comparatively evaluated due to the lack of a sensitive contaminant assay. Here, we applied our ContamSPOT assay to compare various bottom-up proteomics sample preparation methods: S-Trap^15^, SP3^18^, SP4 ^18^, and acetone precipitation^25^, as well as peptide cleanup methods: SP2^17^, Thermo Scientific HiPPR detergent removal resin, and HLB desalting plate. HEK protein lysate containing 2% SDS and 1% Triton was used as the starting material. ContamSPOT, peptide colorimetric assay, and LC-MS were used as readouts. As shown in **Figure 4A, 4B**, acetone precipitation and detergent removal resin cannot sufficiently remove detergents from the sample. Beads-based sample preparation (SP3, SP4, SP2) provided more sufficient detergent removal compared to trap-resin based spin columns (S-Trap). Detergent amount after S-Trap was below 0.01% and did not significantly influence peptide/protein identifications.^10^ We recommend using five or more wash cycles (manufacture recommends 3 times) for both S-Trap and SP4 methods to sufficiently remove detergents to protect chromatography columns and MS instrument. SP3 provided the best overall detergent removal and peptide recovery among all methods (**Figure 4C, 4D**). SP2 cleanup sufficiently removed detergents at the peptide level but had significant sample loss, consistent with previous report.^17^ Principal component analysis showed sample clustering based on sample preparation methods. S-trap, SP3, and SP2 showed better reproducibility compared to SP4 method (**Figure 4E, 4F, Supplemental Figure S3**). But SP4 used glass beads and are much cheaper compared to magnetic beads or S-trap. We also found that even trace amount of detergent could wash away packing materials from reverse phase (C18 or HLB) desalting column, causing serious polymer contaminations in LC-MS. Therefore, if detergents were used in the lysis buffer, ethyl acetate cleanup can be conducted as a routine step before desalting to avoid polymer contamination from desalting columns.

**Figure 4:**
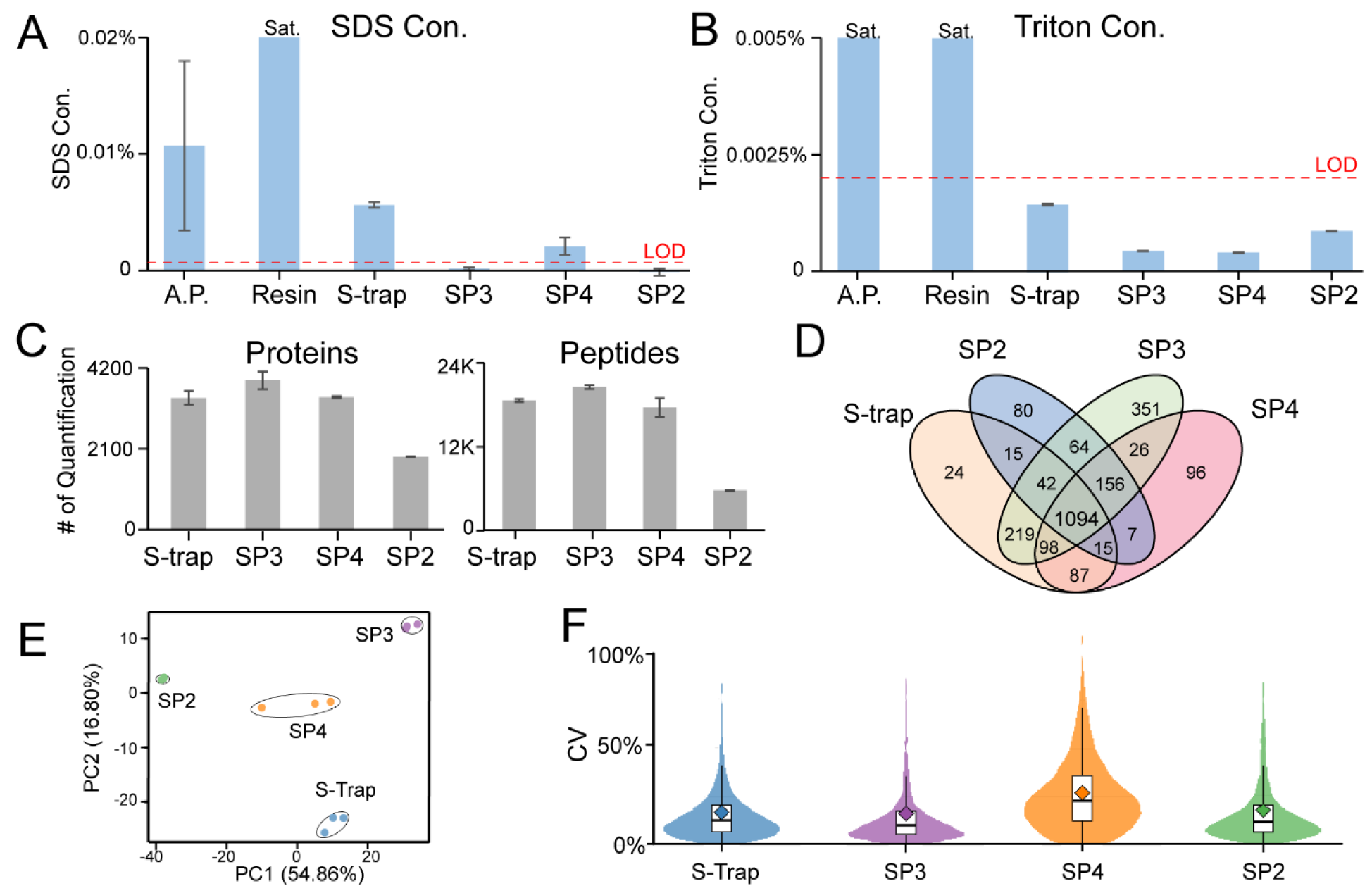
Applying ContamSPOT assays to compare different protein and peptide sample preparation methods. (A) ContamSPOT colorimetric measurement of SDS concentrations after various sample preparation methods. Sat. indicates saturated signal. (B) ContamSPOT fluorescence measurement of Triton concentrations. (C) Number of quantified proteins and peptides from different methods. (D) Venn diagram of reproducibly quantified proteins in different methods. (E) Principal component analysis of different methods with three replicates. (F) Violin plot showing coefficient of variance of different methods.

### A Practical User Guide for ContamSPOT Assay

Given the merits of our ContamSPOT assay (sensitive, rapid, simple, low sample input, and low-cost), we anticipate that ContamSPOT can be a very useful method for the MS community. To assist the use of ContamSPOT in other laboratories, we provided a practical decision tree (**Figure 5**) and a step-by-step protocol with a troubleshooting guide (**Supporting Information**) for users to incorporate ContamSPOT to their routine sample preparation workflow. ContamSPOT should be performed after routine protein digestion and peptide cleanup steps. Researchers should use prior knowledge of possible contaminations in the sample based on the buffers and experimental steps to select ContamSPOT colorimetric or fluorometric assay. ContamSPOT colorimetric assay should be used if there are potential ionic contaminations, such as SDS, TEAB, Tris, AmBC, etc. ContamSPOT fluorometric assay should be used if potential non-ionic contaminations exist, such as Triton, Tween, NP-40, CHAPS, etc. If both ionic and non-ionic reagents were used during sample preparation, we recommend doing colorimetric assay first and then fluorometric assay, which only consumes 1-2 µL of sample. Our ContamSPOT assay can be used for both quantitative and qualitative analyses. A quick qualitative check of sample contamination can use as little as 0.5 µL of sample, but accurate quantification of contaminant concentrations requires at least 1 µL of sample to reduce pipetting variability.

**Figure 5:**
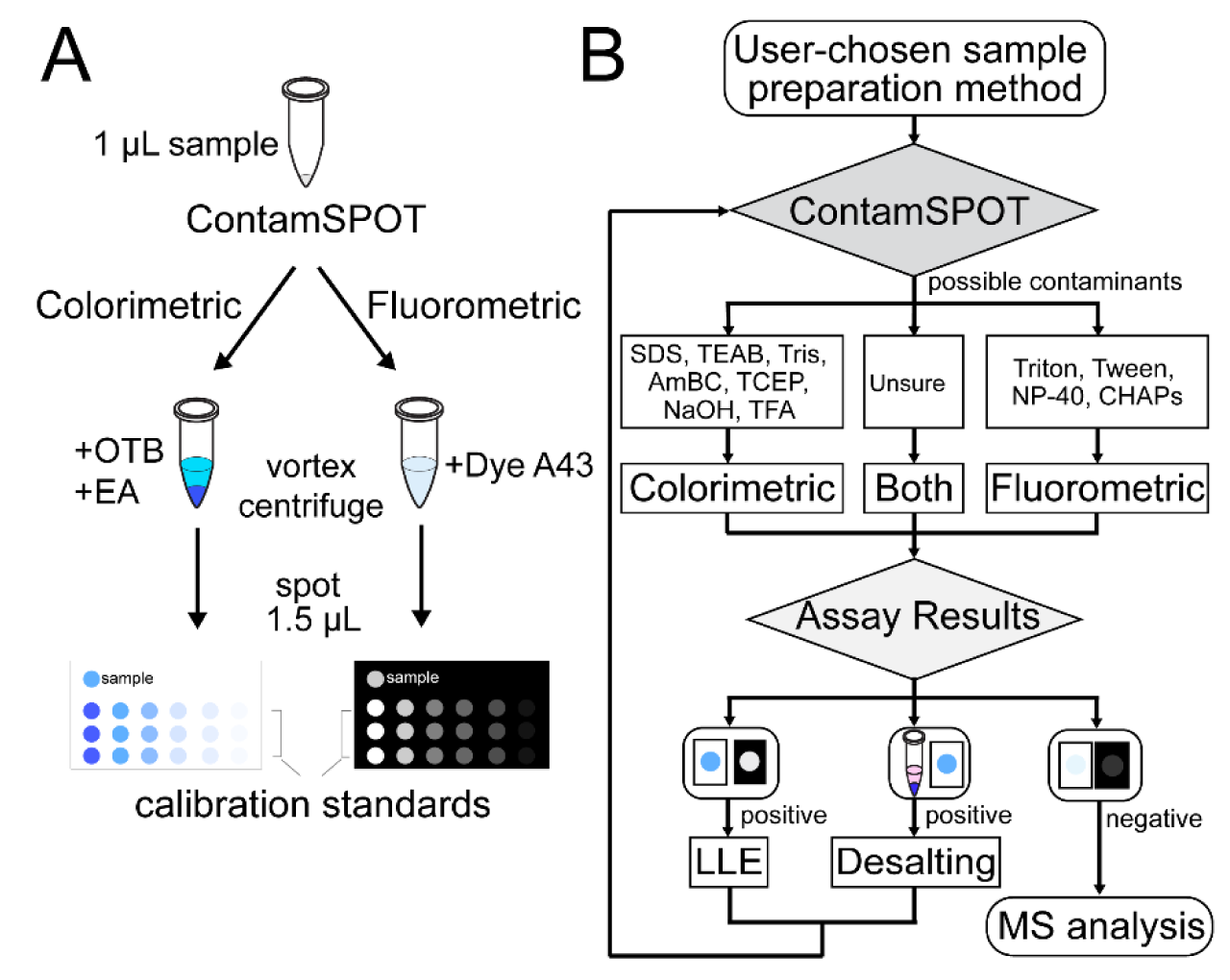
A practical user guide for ContamSPOT. (A) Schematic workflow for ContamSPOT colorimetric and fluorometric assay. (B) Decision tree to implement ContamSPOT to detect, quantify, and remove contaminants prior to MS analysis.

If ContamSPOT showed negative result (i.e., no blue color on TLC plate in colorimetric assay, and/or no signal in fluorometric assay), the samples could safely proceed to MS injection. If ContamSPOT showed positive result, further cleanup of the sample is necessary. Interestingly, the colorimetric assay could distinguish when there are salt contaminations by showing a pink color in the ethyl acetate layer before spotting. We have demonstrated that ethyl acetate cleanup can rapidly and effectively remove detergents, such as SDS, Triton, NP-40, and Tween. However, it is worth noting that ethyl acetate extraction cannot remove CHAPs and salts. While volatile salts (*e.g.* TEAB and AmBC) could be removed by drying down with acidic pH, nonvolatile salts (*e.g.* Tris) will require an additional desalting step. For samples that use detergent in the workflow, water-saturated ethyl acetate extraction can be included as a routine cleanup step to remove possible detergent contamination and then test by ContamSPOT. However, any additional sample preparation step can introduce possible sample loss and variation and therefore should be conducted with caution. We recommend first conducting qualitative ContamSPOT colorimetric/fluorometric test (which takes 1 µL of sample within a few minutes and less than $1.50 per sample to conduct both tests) to decide further cleanup steps (**Figure 5**). The cleaned samples can be tested again with ContamSPOT and proceed to MS analysis when negative results are obtained. ContamSPOT colorimetric and fluorometric assays could also be used to test detergent contamination from various LC-MS experiments for lipid, metabolite, and intact protein analyses (we have not tested these experiments). Ethyl acetate cleanup can only be used for hydrophilic analytes to avoid sample loss into the ethyl acetate layer.

## CONCLUSIONS

In this study, we developed a sensitive, rapid, and low-cost method to quantify and remove contaminants prior to MS analysis, namely ContamSPOT. We developed both colorimetric and fluorometric assays to detect a range of ionic and non-ionic detergents, salts, and chemicals that are commonly used in MS workflows. We tested a total of 22 commonly used chemicals, 11 of which can be quantified by ContamSPOT. Particularly, trace amounts of detergents (*e.g.* 0.0004% SDS, 0.002% Triton) can be detected at nanogram levels to prevent damaging the chromatography and MS instruments. Although ContamSPOT is not selective to each positively tested chemical, we provided performance characteristics that can provide clues for identification. A simple Yes or No answer is sufficient to decide whether the samples are clean enough for MS analysis or should go through additional cleanup steps. Each sample costs less than $1.50 to run both assays making it practical for large scale application. ContamSPOT is also capable of high-throughput sample processing with a multi-channel pipette or a liquid handler. Enabled by ContamSPOT assay as the readout, we optimized ethyl acetate liquid-liquid extraction to rapidly remove trace amounts of detergents from peptide samples with minimal sample loss. We further applied ContamSPOT to compare various proteomic sample preparation workflows, regarding effectiveness to remove contamination, peptide recovery, and reproducibility. We provided a practical guide and a decision tree to help researchers implement ContamSPOT as a rapid quality checkpoint before MS injection. ContamSPOT assay can also be easily adopted when developing or evaluating sample preparation methods as a unique readout for sample cleanness.

## SUPPORTING INFORMATION

Supplemental Step-by-Step Protocol. Supplemental Figures: Effect of emission time of the fluorescence imager on the ContamSPOT fluorometric assay; Evaluation of ethyl acetate cleanup efficiency with 5-fold ethyl acetate volume; LC-MS base peak chromatograms of HEK protein digest before and after ethyl acetate cleanup; Peptide level spearman correlation heat map across different sample preparation methods with three replicates.

## DATA AVAILABILITY

All MS raw files are publicly available on the MassIVE Repository with the identifier MSV000093387. Other data are available in the main text and supporting information.

## AUTHOR CONTRIBUTIONS

1. N. S. and L. H. designed the study. N. S., K. B., and W. M. conducted the experiments. N. S. and
2. H. L. analyzed the data. ContamSPOT assay was tested by all Hao Lab members. N. S., H. L., W. M., and L. H. wrote the manuscript with input from all Hao Lab members.

## ACKNOWLEDGEMENTS

This study is supported by the NSF CAREER grant (CHE 2239214, Hao). L.H. acknowledges the Cottrell Scholar Award. N.S. acknowledges the Bourbon F. Scribner Endowment Fellowship. K. B. acknowledges the NSF Research Experiences for Undergraduates (REU) summer program (CHE 1950193). The authors thank Prof. Joseph Meisel for the access to the Fluorescence Imager.

The authors declare no competing financial interests.

## Notes

### Competing Interest Statement

The authors have declared no competing interest.

