## Supporting Information for "Contaminant Spot Check and Removal Assay (ContamSPOT) for Mass Spectrometry Analysis"

#### Step-by-Step Protocol

Noah Smeriglio, Haorong Li, Wan Nur Atiqah binti Mazli, Katharine Bendel, Ling Hao*

Department of Chemistry, The George Washington University, Science and Engineering Hall 4000, 800 22nd St., NW, Washington, D.C., 20052, USA

*Corresponding author

Ling Hao

Assistant Professor

Department of Chemistry

Department of Biochemistry and Molecular Medicine

George Washington University

**Note: Please cite our publication when using this protocol.**

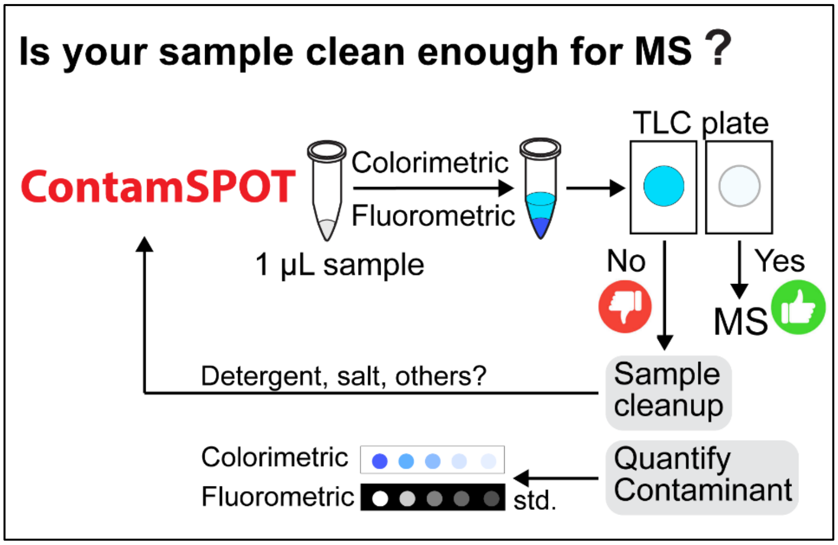

### 1 Introduction of the Method

We aim to provide a sensitive, rapid, simple, and low-cost method to detect, quantify, and remove contaminants before mass spectrometry analysis, namely ContamSPOT. Only one microliter of sample is needed to quantify contaminants by ContamSPOT colorimetric or fluorometric assay on a thin layer chromatography (TLC) plate. ContamSPOT assay can serve as a quick quality checkpoint prior to MS injection, as well as a unique readout of sample cleanness for method development.

This step-by-step protocol provides detailed experimental steps to detect contaminants using colorimetric or fluorometric spot assay as well as ethyl acetate liquid-liquid extraction to remove contaminants.

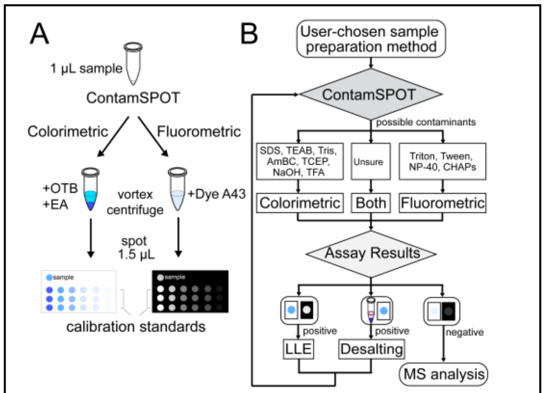

**Figure 5 from the manuscript:** schematic workflow (A) and decision tree (B) to use ContamSPOT.

**Table 1 from the manuscript: Summary of chemicals that can be analyzed by ContamSPOT assay.**

| Chemical | Type | ContamSPOT | $R^2$ | CV | LOD | LOQ | Removal method |
|---|---|---|---|---|---|---|---|
| SDS | Detergent | Colorimetric | 0.991 | 4.6% | 0.0004% | 0.0013% | LLE[a] |
| TEAB | Salt | Colorimetric | 0.98 | 3.1% | 0.9 mM | 2.7 mM | Desalting[b] |
| AmBC | Salt | Colorimetric | 0.99 | 1.5% | 3 mM | 8 mM | Drying/desalting |
| Tris | Salt | Colorimetric | 0.995 | 5.4% | 4 mM | 12 mM | Desalting |
| TCEP | Reducing agent | Colorimetric | 0.99 | 7.5% | 8 mM | 24 mM | Desalting |
| NaOH | Base | Colorimetric | 0.97 | 0.4% | 0.11 mM | 0.34 mM | Desalting |
| TFA | Acid | Colorimetric | 0.99 | 6.6% | 0.0007% | 0.002% | Drying/LLE |
| Triton X-100 | Detergent | Fluorometric | 0.95 | 2.7% | 0.002% | 0.005% | LLE |
| Tween-20 | Detergent | Fluorometric | 0.98 | 4.5% | 0.02% | 0.06% | LLE |
| NP-40 | Detergent | Fluorometric | 0.96 | 2.9% | 0.02% | 0.05% | LLE |
| CHAPS | Detergent | Fluorometric | 0.99 | 4.5% | 0.02% | 0.06% | Cannot be removed by LLE |
| Others[c] | | - | - | - | - | - | - |

[a]Liquid-liquid extraction using ethyl acetate.
[b]Desalting with reverse phase C18 or HLB cartridges/plates.
[c]Tested negative in ContamSPOT assay, including NaCl, urea, iodoacetamide, acrylamide, dithiothreitol, formic acid, hydrochloric acid, EDTA, acetone, acetonitrile, and methanol.

#### 2 ContamSpot Colorimetric Assay Protocol

1. Add 1 µL of 0.1% o-toluidine blue (OTB) dye to a 0.2 ml tube.
   (Note: tube size and pipetting accuracy influence the reproducibility of the assay.)
2. Add 1 µL of sample to the tube.
   (Note: for quantification, a series concentration of contaminant standard need to be used.)
3. Add 3 µL of HPLC-grade ethyl acetate (EA) to the tube.
4. Vortex the tube for 15 s followed by 15 s of centrifugation using a benchtop mini-centrifuge.
5. Spot 1.5 µL of the ethyl acetate layer on a thin-layer chromatography (TLC) plate.
   (Note: If sample volume is not limited, can use 2 µL of sample, 2 µL of OTB, and 5 µL of EA, and spot 2 µL. If sample volume is limited, can use 0.5 µL of sample for a qualitative test.)
6. Wait for the spot to completely dry.
   (Note: Spotting is stable for ~1 h before the color begins to fade. For large sample sets, we recommend closing each tube cap to avoid solvent drying out before spotting.)
7. Take a photo (with flash-light). Qualitative Yes or No can be directly visualized by eye as a blue color. Detergent standards (>1.5X LOD) and buffer control can be used as references for comparison. Quantitative analysis requires further image analysis by ImageJ or other software.

#### 3 ContamSPOT Fluorometric Assay Protocol

1. Dilute ProFoldin's 100x Dye A43 with Reagent 1 to 1x Dye A43 using 10 µL with 990 µL of Reagent 1 from the ProFoldin Assay Kit.
2. Dilute 100x Reagent 2 using 10 µL into 990 µL of HPLC grade water.
3. Add 1 µL of sample to the tube.
   (Note: for quantification, a series concentration of contaminant standard need to be used.)
4. Add 2 µL of 1x Dye A43 to each tube.
5. Add 1 µL of 1x Reagent 2 to each tube.
6. Vortex each tube for 5 s followed by 5 s of centrifugation.
7. Wait for 5 min for sufficient reaction.
8. Spot 2 µL of the mixture on a TLC plate.
9. Image by a Fluorescence Imager with 535 nm wavelength emission.

### 4 ImageJ Analysis

##### 4.1 Colorimetric Image Processing

1. Following imaging of the TLC plate, upload the photo to ImageJ using File →Open.
2. Crop the background of the plate by the box selection function followed by Image → Crop.
3. Subtract the background with a rolling ball radius of 50 pixels and light background selected.

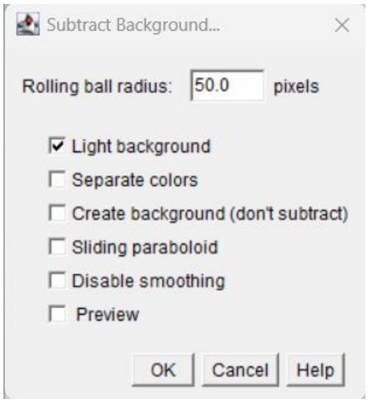

4. Double click the oval brush selection and check the enable selection brush function and set pixels to the desired value. (Note: Pixel sizes can vary based on image quality and spot size. Use a consistent number for each assay.)

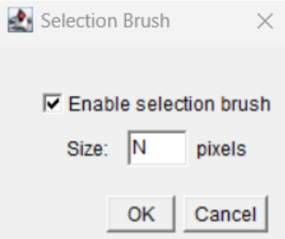

5. To select a spot for measurement, click the center of the spot and a ringed selection should cover the spot size.
6. The spot should be exactly covered which requires consistent spotting every time. To analyze the spot, click Control + M. This will generate a mean and integrated density. To de-select and move onto the next spot, click Control + Shift + A.
7. Analyze each spot with the same pixel size, if the spot is deformed still collect the data as a concentration of signal in one area will be accounted for when evaluating the integrated density.

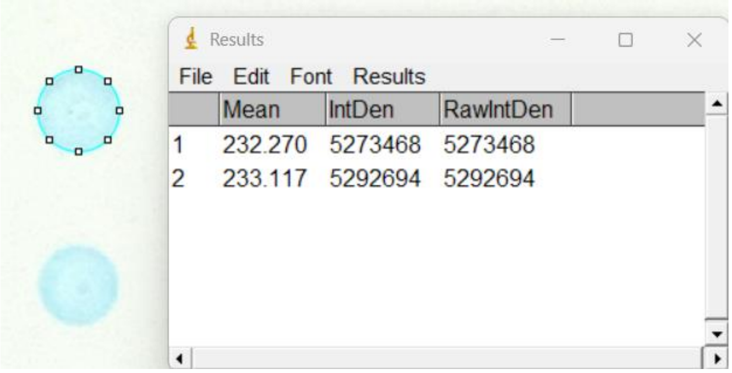

8. Integrated densities of all sample spots should be lower than the control. A relative signal intensity can be calculated using below equation:

$$Relative\ Signal\ Intensity = Sample - Control$$

**4.2 Fluorometric Image Processing**

1. Following imaging of the TLC plate, upload the photo to ImageJ using File →Open.
2. Crop the image to focus on areas spotted during the fluorescent spotting step by selecting the area with the box selection tool followed by Image → Crop.
3. Subtract the background: Process → Subtract Background with a rolling ball radius of 50 pixels.
4. Select the oval selection brush followed by selecting an appropriate pixel size for the spots.

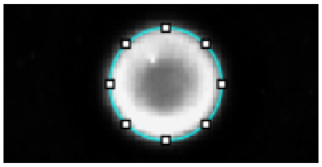

5. Enter Control + M for integrated density collection and a results table will automatically pop up.
9. On the image, enter Control + Shift + A to deselect that circle, repeat for all spots then copy all results to Excel. A relative signal intensity can be calculated using below equation:

$$Relative\ Signal\ Intensity = Sample - Control$$

### 5 Ethyl Acetate Liquid-Liquid Extraction

##### 5.1 Water Saturated Ethyl Acetate

1. In a glass bottle, add 10 mL of HPLC grade water to 50 mL of HPLC grade ethyl acetate and shake vigorously for at least 1 minute.
2. Allow the bottle to settle. A water layer should settle at the bottom below ethyl acetate layer.

##### 5.2 Liquid-Liquid Extraction of Detergents

1. Add 10x sample volume of water-saturated ethyl acetate to the sample.
   (Note: If sample volume is higher than 100 µL, can concentrate below 100 µL before adding ethyl acetate to reduce the total volume used and fit in a 2 mL tube.)
2. Vortex the sample tube for 30 s followed by 30 s of centrifugation on a benchtop mini centrifuge.
3. Allow the water to settle below the organic layer (10 s) then carefully remove the ethyl acetate layer, leaving a small amount of organic layer to ensure no water/sample layer is removed.
4. Repeat the wash cycle for at least 3 times.
5. Dry down the sample to completely remove ethyl acetate before MS injection.

##### 5.3 Testing for Sample cleanness during LLE

(Can be conducted during LLE to check sample cleanness without consuming any sample)

1. During LLE washing cycles, transfer the most recent ethyl acetate wash layer to a new tube and dry it down.
2. Reconstitute with minimal volume of water (5-20 µL) and vortex for complete dissolution.
3. Conduct the qualitative ContamSPOT colorimetric or fluorometric assay to test for contaminations.
4. If ContamSPOT showed negative results, LLE is complete. If ContamSPOT showed positive results, conduct more ethyl acetate washes.

### 6 Troubleshooting Guide

#### 6.1 Colorimetric and Fluorometric Assay Troubleshooting

| Issue: | Potential Reason(s): | Action: |
|---|---|---|
| Signal of standards are faint in Colorimetric assay | The standards may have degraded over time | Make fresh standards |
| Signal of standards are faint in Fluorometric assay | The extraction<br>OR<br>Emission time is not long enough. | Wait longer before spotting the mixture.<br>OR<br>Turn off automated emission data collection and manually increase the emission time. |
| Extremely strong signal | Accidental pipetting of the aqueous layer.<br>OR<br>Sample is extremely contaminated. | Angle the tube to ensure no dark blue droplets in the organic layer when taking the liquid for spotting.<br>OR<br>Cleanup the sample. |
| High variation of quantification | Pipetting or spotting variation. | Quantitative assay requires at least 1 uL of sample. If sample volume is not limited, increase to 2 µL of sample volume. |
| No layer formed in colorimetric assay | Likely one of the 3 reagents was not added.<br>OR<br>Sample is contaminated by organic solvent. | Repeat that sample/standard.<br>OR<br>Dry down samples and redissolve in water. |

##### 6.2 Liquid-Liquid Extraction Troubleshooting

| Issue: | Potential Reasons: | Action: |
| --- | --- | --- |
| Not forming layers | Sample containing high amount of organic solvent | Dry down samples and redissolve in water. |
| A solid precipitate layer formed between the ethyl acetate and water. | High salt content or proteins, high urea concentrations can cause a salt line. | If samples are lysed in urea with the absence of detergent entirely, conduct desalting then test ContamSPOT. |
| Post LLE clean-up still show positive results of ContamSPOT | Not enough wash cycles<br>OR<br>The sample is contaminated with chemicals that cannot be removed via LLE (salt) | Increase number of ethyl acetate washes.<br>OR<br>Check the ethyl acetate layer prior to spotting. If the layer is pink, desalt the sample. Volatile salts can be removed by drying down in acidic pH (adding formic acid) |
| Significant sample loss after LLE | Volume of the samples are too high. | Concentrate the samples to below 100 µL. Use water-saturated EA instead of pure EA. |

### 7 List of Reagents

| Reagent | Purpose | Vendor |
|---|---|---|
| Ethyl Acetate (EA, ACS grade) | Colorimetric Assay | Fisher Scientific |
| 1% O-Toluidine Blue (OTB, Laboratory grade) | Colorimetric Assay | Carolina Biological |
| Thin line chromatography (TLC) plate | ContamSPOT | Sorbtech |
| Detergent Assay Kit DAK100 | Fluorometric Assay | ProFoldin |
| Ethyl acetate (EA, HPLC grade) | Solvent | Sigma Aldrich |
| Trifluoroacetic Acid (TFA, Optima) | Acid | Fisher Scientific |
| Formic Acid (FA, Optima) | Acid | Fisher Scientific |
| Sodium dodecyl sulfate (SDS, 20%) | Detergent | Fisher Scientific |
| NP-40 | Detergent | Millipore Sigma |
| Triton X-100 | Detergent | Thermo Fisher Scientific |
| TWEEN® 20 | Detergent | Sigma Aldrich |
| CHAPS | Detergent | Fisher Scientific |
| Triethyl Ammonium Bicarbonate buffer (TEAB) | Lysis Salt Buffer | Millipore Sigma |
| Ammonium Bicarbonate (AmBC) | Lysis Salt Buffer | Thermo Fisher Scientific |
| Tris Base | Lysis Salt Buffer | Fisher Scientific |
