## Supplemental Figure for "Contaminant Spot Check and Removal Assay (ContamSPOT) for Mass Spectrometry Analysis"

#### Supporting Information

Noah Smeriglio, Haorong Li, Wan Nur Atiqah binti Mazli, Katharine Bendel, Ling Hao*

Department of Chemistry, The George Washington University, Science and Engineering Hall 4000, 800 22nd St., NW, Washington, D.C., 20052, USA

*Corresponding author

Ling Hao

Assistant Professor

Department of Chemistry

Department of Biochemistry and Molecular Medicine

George Washington University

### Table of Contents


- Supplemental Figure S1: Effect of emission time of the fluorescence imager on the ContamSPOT fluorometric assay.

- Supplemental Figure S2: Evaluation of ethyl acetate cleanup efficiency with 5-fold ethyl acetate volume.

- Supplemental Figure S3. LC-MS base peak chromatograms of HEK protein digest before and after ethyl acetate cleanup.

- Supplemental Figure S4: Peptide level spearman correlation heat map across different sample preparation methods with three replicates.

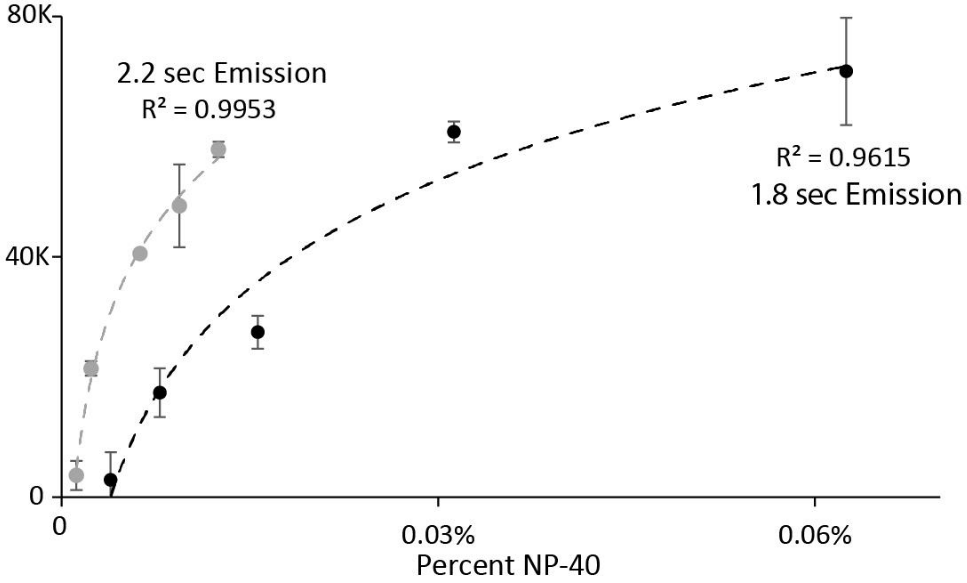


**Supplemental Figure S1: Effect of emission time of the fluorescence imager on the ContamSPOT fluorometric assay.** Emission time needs to be optimized for each Fluorescence Imager to obtain non-saturated and quantifiable signals for various concentrations of detergents.

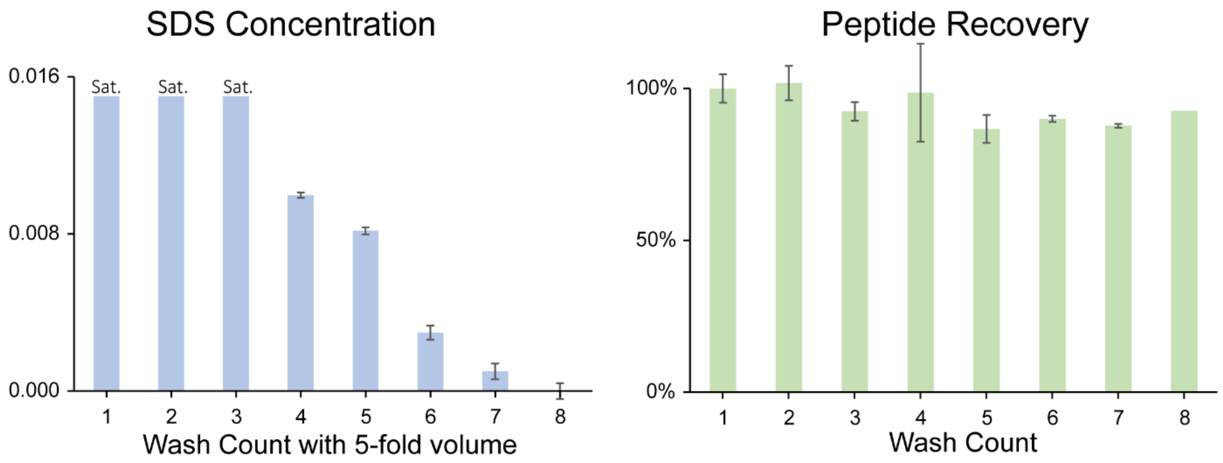


**Supplemental Figure S2: Evaluation of ethyl acetate cleanup efficiency with 5-fold ethyl acetate volume.** (A) SDS concentrations after multiple washes. 0.25% SDS was spiked into peptide samples as the starting material. (B) Peptide recovery after multiple washes. If 5-fold ethyl acetate volume (ratio to sample volume) was used, eight washes are needed to completely remove SDS without causing significant sample loss.

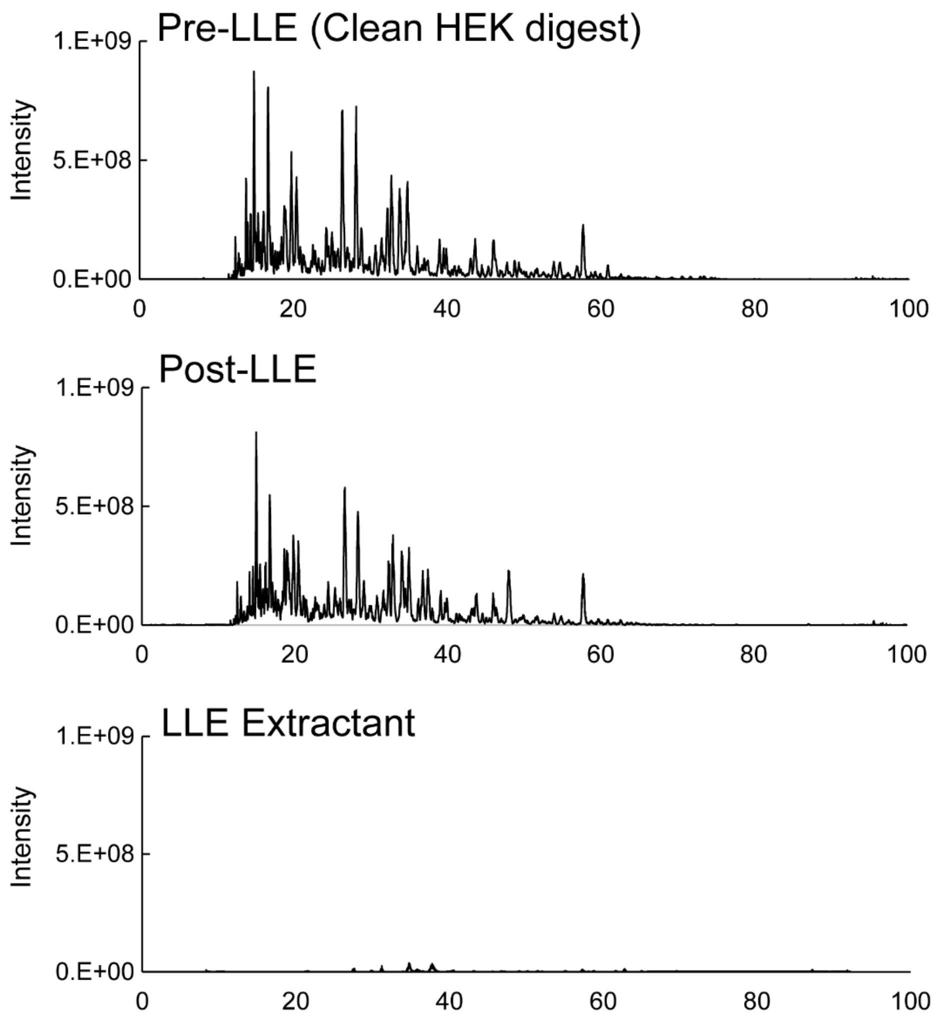


**Supplemental Figure S3. LC-MS base peak chromatograms of HEK protein digest before and after ethyl acetate cleanup.** Minimal signal loss was observed after LLE. LLE extract has no peptide signals. ACS grade ethyl acetate generated several singly charged contaminant peaks in LLE extractant. Therefore, HPLC grade ethyl acetate should be used.

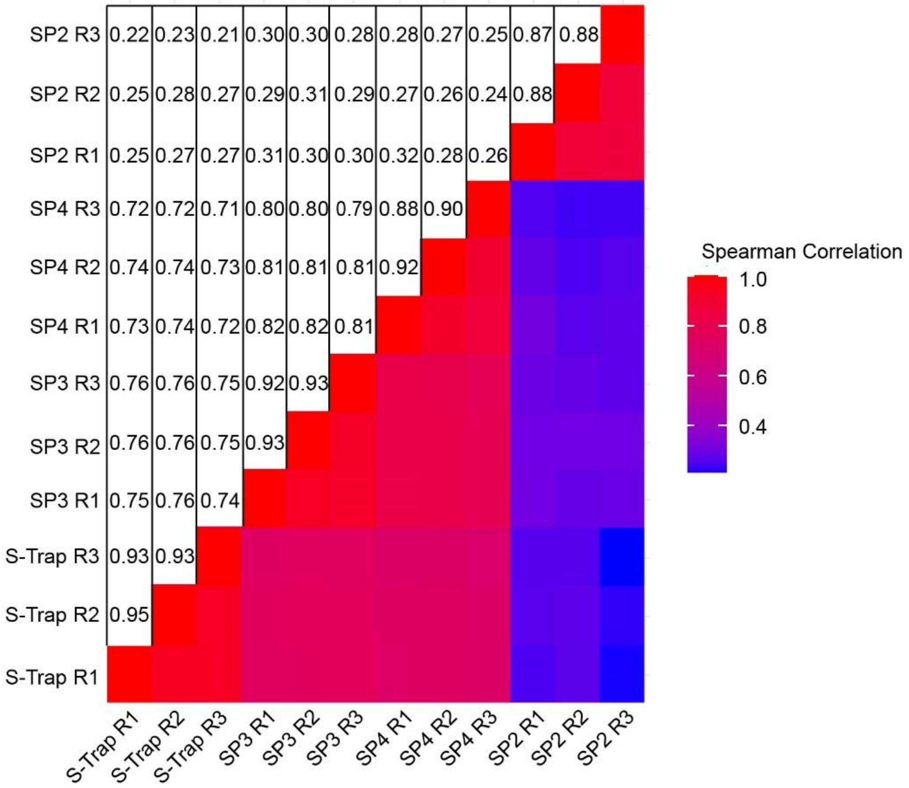


**Supplemental Figure S4: Peptide level spearman correlation heat map across different sample preparation methods with three replicates.** Replicates from each method showed strong correlation and tight clustering. SP2 has the lowest correlation among all methods because of significant sample loss.
